# Inhibition of the Notch signaling pathway promotes AQP2 plasma membrane accumulation in renal epithelial cells by depolymerizing actin and reducing endocytosis

**DOI:** 10.64898/2026.08.11.744289

**Authors:** Asma Tchakal-Mesbahi, Huihui Huang, Jake Ross, Richard Bouley, Dennis Brown

## Abstract

The Notch signaling pathway plays a central role in development and cell fate determination. Its function depends on tightly regulated intracellular trafficking of the Notch receptor and the Notch intracellular domain (NICD) after cleavage by γ-secretase. Notch signaling is essential for principal cell differentiation within the renal collecting duct and for proximal-distal patterning during kidney development. Notch activity has also been shown to influence the trafficking of several membrane proteins, including nephrin in kidney cells and monocarboxylate transporter 1 in brain endothelial cells. Aquaporin-2 (AQP2) is the key vasopressin-regulated water channel in the collecting duct, and proper AQP2 trafficking and recycling are required for physiologically appropriate urine concentration. To determine whether and, if so, how Notch signaling modulates AQP2 trafficking, we performed studies using LLCPK1 renal epithelial cells stably expressing AQP2 (LLCPK1-AQP2). Exposing cells to 35 μM DAPT (which inhibits y-secretase, preventing cleavage and activation of Notch receptor signaling) for 30 min significantly increased AQP2 membrane accumulation in LLCPK1-AQP2 cells as revealed by immunofluorescence staining. Using a rhodamine-transferrin internalization assay, we found that DAPT reduced clathrin-mediated endocytosis by 60%. This blockade increases AQP2 membrane accumulation by preventing the reinternalization of AQP2 that is delivered to the plasma membrane by exocytosis during its constitutive recycling pathway. Using an F-actin polymerization assay, we then found that Notch inhibition decreases F-actin polymerization by de-activating the small GTPase RhoA, using GSTRBD, a substrate that binds to active RhoA, as seen by western blotting using phospho-specific antibodies. Because actin polymerization is required for AQP2 endocytosis, RhoA inhibition by DAPT would result in the decreased internalization of AQP2 that we observed by immunofluorescence. While the mechanism by which DAPT inhibits RhoA activity remains to be determined, our study shows that AQP2 trafficking is regulated by the Notch signaling pathway in vitro and suggests that modulation of Notch signaling may represent a novel strategy to address water balance disorders that involve defects in the AQP2 trafficking process.

## Introduction

The water channel aquaporin-2 (AQP2) plays an essential role in maintaining fluid balance by regulating water reabsorption in the renal collecting duct (CD) (Knepper et al., 2015; Nielsen et al., 2002). Within principal cells, AQP2 is predominantly located in intracellular vesicles located near the apical surface and accumulates in the plasma membrane in response to physiological stimuli that increase urinary concentration (Brown, 2003; Nielsen et al., 2002). This trafficking is primary regulated by the antidiuretic hormone, vasopressin (VP) in response to hypovolemia or high blood osmolality, initiating a cascade of signaling as a result of its binding to the vasopressin type 2 receptor (V2R). This involves activation of adenylyl cyclase, generation of cyclic AMP, and stimulation of protein kinase A (Brown & Fenton, 2015; Fenton & Moeller, 2008; Knepper & Inoue, 1997; Knepper et al., 2015; Ranieri et al., 2019). The resulting phosphorylation of AQP2 at serine 256 in the C-terminal region of AQP2 is both required and sufficient to cause AQP2 accumulation into the apical membrane and enhance principal cell/collecting duct water permeability (Fushimi et al., 1997; Katsura et al., 1997).

Several clinical disorders are closely linked to the impairment or improper regulation of AQP2 trafficking. Mutations affecting either AQP2 or V2R are responsible for most cases of congenital and acquired nephrogenic diabetes insipidus, a condition characterized by impaired urinary concentrating (Bichet, 2006; Radin et al., 2012; Sands et al., 2006). Conversely, excessive AQP2 membrane expression contributes to pathological fluid retention observed in conditions such as congestive heart failure, pregnancy-associated edema, obstructive uropathy, and lithium-induced renal dysfunction (Knepper et al., 2015; Kortenoeven & Fenton, 2014).

The Notch signaling pathway is a highly conserved mechanism of intercellular communication that influences numerous developmental and physiological processes. In mammals, this pathway consists of four receptors (Notch1 through Notch4) and five ligands, including members of the Delta-like and Jagged families (Artavanis-Tsakonas et al., 1995; Bray, 2016; Kopan & Ilagan, 2009). Activation occurs when ligand binding induces sequential proteolytic cleavage of the receptor, ultimately releasing the Notch intracellular domain. This intracellular fragment translocates to the nucleus, where it functions as a transcriptional regulator controlling genes involved in cell differentiation, survival, proliferation, and lineage specification (Iso et al., 2003; Kopan & Ilagan, 2009). During renal organogenesis, Notch signaling plays a central role in nephron patterning and epithelial cell fate determination. Both Notch1 and Notch2 are expressed in developing glomerular and tubular structures and contribute to the specification of proximal nephron segments and podocyte differentiation (Cheng et al., 2003; McCright et al., 2002). Among these receptors, Notch2 appears particularly critical, as its loss during early kidney development results in failure of nephron formation (McCright et al., 2002). Although expression of Notch signaling components declines in mature kidneys, accumulating evidence indicates that this pathway is reactivated during renal injury and repair. Aberrant Notch activation has been implicated in a variety of kidney pathologies, including acute kidney injury and chronic kidney disease, where it contributes to fibrotic remodeling (Bielesz et al., 2010; Niranjan et al., 2008). Experimental inhibition of Notch signaling reduces fibrosis in animal models, and targeted disruption of downstream transcriptional mediators decreases expression of profibrotic markers (Bielesz et al., 2010; Kramer et al., 2016; Soni et al., 2019; Xiao et al., 2014). Furthermore, Notch activation in glomerular epithelial cells has been associated with proteinuria and progressive glomerular scarring (Niranjan et al., 2008).

However, more recent studies suggest that Notch signaling influences cellular functions beyond its “canonical” role in development and transcriptional regulation, including an involvement in membrane protein trafficking. For example, activation of Notch signaling in podocytes promotes endocytosis of nephrin through dynamin-dependent mechanisms, leading to decreased surface expression and structural injury (Waters et al., 2012). Additionally, Notch signaling interacts with Wnt/β-catenin (Ando et al., 2016) pathways to regulate trafficking of metabolic transporters such as monocarboxylate transporter-1 in the blood-brain barrier (Liu et al., 2016). Importantly, a role of Wnt5a signaling in regulating AQP2 trafficking independently of the PKA/cAMP pathway and AQP2 phosphorylation has been reported (Ando et al., 2016).

Increasing evidence has also linked NOTCH signaling to cytoskeletal remodeling and many studies have shown that Notch receptors reorganize actin filaments in several cell types. For instance, Notch 3 affects vascular contractility by modulating cytoskeletal structure in resistance arteries and alters actin architecture in vascular smooth muscle cells (Domenga et al., 2004). Similarly, Notch1 signaling limits invasive behavior in breast cancer cells by suppressing actin polymerization (Domenga et al., 2004; Rizzo et al., 2013; Sahlgren et al., 2008). These findings are particularly relevant to collecting duct physiology because cytoskeletal dynamics play a key role in controlling AQP2 trafficking (Noda et al., 2004; Sasaki et al., 2014; Tamma et al., 2001; Yui et al., 2012).

The actin cytoskeleton is an essential component of clathrin-mediated endocytosis, providing mechanical support for the formation and internalization of clathrin-coated vesicles (Ridley, 2006; Ridley et al., 2003). Since this endocytic pathway is a major determinant of AQP2 retrieval from the apical membrane, inhibition of clathrin-dependent internalization leads to increased membrane retention of AQP2 (Lu et al., 2004; Sun et al., 2002). RhoA, a small GTP-binding protein, is a major regulator of actin filament assembly and functions as a critical mediator of vesicular transport processes (Ridley, 2006; Ridley et al., 2003; Tamma et al., 2001). Experimental suppression of RhoA activity or disruption of actin polymerization enhances the surface accumulation of AQP2 (Klussmann et al., 2001; Li et al., 2011; Tamma et al., 2003). Importantly, Notch signaling can modulate RhoA activity and thereby influence cytoskeletal organization (Liu et al., 2019; Waters et al., 2012; Xiao et al., 2014).

The small molecule DAPT (N-[(3,5-Difluorophenyl) acetyl]-L-alanyl-2-phenyl]glycine-1,1-dimethylethyl ester) is a powerful tool to examine the Notch signaling pathway (Ba et al., 2012; Boucher et al., 2020; Li et al., 2017; Wu & Kitajewski, 2009; Zhang et al., 2019). It inhibits γ-secretase activity, thereby preventing Notch receptor enzymatic hydrolysis, which blocks intracellular domain release and transcriptional activation in the Notch pathway (Hans et al., 2020; Yin et al., 2020). Importantly, DAPT is reported to have little or no effect on other cellular signaling pathways (Sun et al., 2019). Based on these considerations, we hypothesized that Notch signaling might regulate actin-dependent water channel trafficking in renal collecting duct cells. We propose that activation of the Notch pathway controls AQP2 membrane targeting by modulating RhoA-dependent cytoskeletal dynamics, thereby influencing water transport and renal concentrating ability.

## MATERIAL AND METHODS

### Chemicals, reagents and antibodies

Lysine vasopressin (LVP), arginine VP (AVP), forskolin (FK), methyl-β-cyclodextrin (MβCD), Notch inhibitor (N-[(3,5-Difluorophenyl)acetyl]-L-alanyl-2-phenyl]glycine-1,1-dimethylethyl ester DAPT) (D5942-5MG) were purchased from Sigma (St. Louis MO). LVP and AVP were used at 20 nM final concentration, FK was used at 10 µM final concentration whereas DAPT was used at a final concentration of 35 μM and MβCD was used at 10 mM final concentration, all in serum free medium. Latrunculin-A which disrupts microfilament organization was purchased from CalBiochem (Billerica MA). Cell culture reagents including medium were purchased from Corning/Cellgro (Manassas VA), fetal bovine serum was purchased from Gibco Brl (Grand Island NY), phosphate-buffered saline (PBS; 10mM sodium phosphate buffer containing 0.9% NaCl, pH 7.4) was obtained from Boston Bioproducts (Boston MA). Alexa Fluor® 555 Phalloidin, GAPDH polyclonal rabbit antibody, rabbit polyclonal antibody to cleaved Notch1 were purchased from Cell Signaling Technology Inc. (Danvers MA). Tetramethylrhodamine transferrin was obtained from Life Technologies (Carlsbad CA). AQP2 polyclonal goat antibody, RhoA mouse polyclonal antibody, donkey anti-rabbit and donkey anti-mouse secondary antibodies coupled to peroxidase were purchased from Santa Cruz Biotechnology (Santa Cruz CA). AQP2 polyclonal rabbit antibody was purchased from Alomone Labs (Jerusalem, Israel). Monoclonal antibody against c-myc was generated from the 9E10 hybridoma cell line, purchased from American Type Culture Collection (ATCC, Manassas VA). Secondary antibodies tagged with Cy3 or FITC were obtained from Jackson Immuno-research Laboratories (West Grove PA); a secondary anti–mouse IgG conjugated to Alexa-488 was from Invitrogen (Waltham, MA).

### Cell culture

LLC-PK1 cells stably expressing wild-type c-myc-tagged aquaporin-2 (LLC-AQP2 cells) (Katsura et al., 1995), LLC-AQP2 cells stably expressing secreted soluble yellow fluorescent protein (LLC-AQP2-ssYFP cells) (Nunes et al., 2008), LLC-AQP2 cells stably expressing a c-myc-tagged alanine 256 mutant c-myc-tagged aquaporin-2 (LLC-AQP2 S256A) (Nunes et al., 2008; Rice et al., 2012) were all cultured at 37°C with a 5% CO_2_ atmosphere in DMEM + 10% FBS. Cells were grown either on coverslips (Electron Microscopy Sciences, Hatfield, PA) or on standard P6 polystyrene Falcon culture dishes (Corning Live Science, Corning, NY) to 70 - 80% confluence prior to performing the experiments.

### Immunofluorescence staining of cells

In initial experiments, cells grown as above were treated with lysine vasopressin (LVP) (10 nm, 15 min) or the NOTCH inhibitor DAPT (DAPT: 35 µM, 30 min) (D5942-5MG; Sigma, St. Louis MO), then fixed in 4% PFA (PFA, Electron Microscopy Sciences, Hatfield, PA) in PBS for 20 min. Fixed coverslips were permeabilized in 0.02% Triton X-100/PBS for 4 min, washed three times with PBS, and blocked in 1% BSA/PBS for 30 min. The coverslips were incubated with primary antibodies for 1 h at room temperature or overnight at 4°C. After washing with PBS three times, coverslips were incubated in secondary antibody for 45 min at room temperature, and after 3 more washes with PBS the coverslips were mounted with Vectashield mounting medium with DAPI. They were examined and imaged using a Nikon 80i microscope (Nikon Instruments, Melville, NY). Image J software was used for image analysis (NIH, http://rsb.info.nih.gov/ij/, Bethesda, MD). Background corrections and contrast/brightness enhancement were performed identically for all images in the same experiment. AQP2 was detected in cells using undiluted culture medium containing mouse anti-c-myc IgG produced by the hybridoma cells followed by a secondary anti–mouse IgG conjugated to Alexa-488.

### Quantification of AQP2 at the plasma membrane

After AQP2 labeling, cell membranes were stained with Alexa 647-conjugated wheat germ agglutinin (Alexa 647 WGA, ThermoFisher Scientific, 0.4 μg/mL) for 10 minutes. After WGA staining, coverslips were washed three times in PBS and mounted on glass slides with VectaShield Plus with DAPI to stain the nucleus (Vector Laboratories, Newark, CA). Quantification was performed using Volocity software (PerkinElmer, Waltham, MA). Under blinded conditions, three regions of interest (ROIs) were defined. First, the nuclear region was determined using only the DAPI channel. Next, with the AQP2 channel turned off, the cytoplasm and membrane regions were defined using the WGA-conjugated Alexa 647 signal to label the plasma membrane. The fluorescence of the AQP2 channel was then evaluated in each predetermined ROI. The membrane fraction of AQP2 was quantified by dividing the intensity in the membrane ROI by the sum of the membrane ROI and cytoplasm ROIs. With this technique, the limit of resolution is such that we cannot distinguish between the plasma membrane and vesicles immediately adjacent to the plasma membrane.

### Endocytosis and exocytosis assays

The endocytosis assay was performed using the following protocol: after 1 h in serum-free DMEM, cells were treated with drugs. Dialysed Texas Red dextran was added to a final concentration of 1.5 mg/ml 10 min before the end of the treatment. Cells were then washed quickly with cold PBS 3 times PBS, then lysed in 150 μl of cold RIPA buffer and protease inhibitor cocktail. Fluorescence signals of the supernatants after centrifugation at 17,000 g for 10 min at 4°C (100 μl) were read on a DTX880 Multimode plate reader (Cheung et al., 2019).

Endocytosis was also monitored using a rhodamine-tagged transferrin ligand (Rho-Tf; Invitrogen). LLC-AQP2 cells were incubated with drugs for 30 min on coverslips placed in 12-well cell culture plates (Corning Live Science, Corning, NY), and Rho-Tf was added to cells to reach a final concentration of 25 µg/ml in 1% BSA / phenol red-free DMEM for 20 min at 37°C. Cells were then washed with cold PBS twice before fixing in 4% PFA in PBS, washed again, and mounted with Vectashield mounting medium with the DNA stain DAPI.

As a positive control, we added the cholesterol-chelating drug methyl-β-cyclodextrin (MβCD), to the cells. At a concentration of 10 mM, MβCD rapidly inhibits most endocytotic events in cells (5-15 mins), as we have previously shown (Cheung et al., 2023; Lu et al., 2004). The coverslips were examined by widefield fluorescence microscopy (Nikon 80i microscope) and analyzed using ImageJ software. The mean fluorescence intensity was measured for each outlined region of interest (ROI), normalized to background fluorescence (of the nucleus), and compared across conditions to assess differences in staining patterns. Background corrections and contrast/brightness enhancement were performed identically for all images in the same experiment (Mamuya et al., 2016).

For the exocytosis assay, we used LLC-AQP2-ssYFP cells. After being cultured in 12 well plates as above, we treated them with either LVP or DAPT as detailed below, and then measured exocytosis of ssYFP secreted into the culture medium by quantifying the fluorescence in the culture medium. The specific protocol was previously described in detail (Nunes et al., 2008) but briefly, the cells were starved by incubation in HBBS (Hank’s buffered saline solution) containing 20 mM HEPES, 25 mM sodium bicarbonate and 11 mM glucose for 1 h, followed by addition of LVP (10 nM) or DAPT (35 µM) for 30 min. After each treatment, 150 µl of medium from each well was transferred to a black half-area 96-well plate and analyzed using a multimode plate reader (model DTX880, Beckman-Coulter, Fullerton, CA). Fluorescence values represent three independent experiments performed in triplicate. Each fluorescence value is reported as a ratio of each background- and zero subtracted value.

### Western blotting

For cell lysate preparation, cells were grown on P6 plates and treated with different drugs as described above, then rinsed 3 times with ice-cold PBS and lysed with a cell scraper in 150 μl lysis buffer containing 8 mM NaF, 0.5% NP-40, 4 mM Na3V04, 0.1% Triton X-100, 0.03% NaN_3_ and 1% protease inhibitor cocktail from Biotool.com (Houston, TX). Samples were incubated on ice, passed at least 5 times through a 27G×1/2 syringe and then centrifuged at 17,000 g at 4°C for 10 min to remove cell debris. The protein concentration in the lysate was measured with the Pierce® BCA Protein Assay Kit from Pierce Biotechnology (Rockford, IL). After adding 3x lithium dodecyl sulfate (LDS) sample buffer (Life Technologies, Carlsbad, CA) the samples were used for western blotting as described previously (Rice et al., 2012). AQP2 was detected using a goat polyclonal antibody, sc9882 (1:1000 dilution) from Santa Cruz Biotechnology (Dallas, TX), and mouse anti GAPDH monoclonal antibody AM4300 (at 1:10000) was from Ambion/Thermo Fisher Scientific, Waltham, MA) and detected with the appropriate HRP-conjugated secondary reagents (as listed in the supplementary table).

### RhoA activity Assay

After being cultured in standard P10 polystyrene Falcon culture dishes to 70-80% confluence, LLC-AQP2 cells were washed with ice-cold PBS, and harvested in lysis buffer with a cell scraper. Equivalent protein amounts of lysates (300-800 µg) were then incubated with 50 µg Rhotekin-RBD beads (Cytoskeleton Inc., Denver, CO) at 4°C on a rotator for 1 hour. The beads were pelleted by centrifugation at 5,000g then washed by removing the supernatant. Then, beads were suspended in 1x lithium dodecyl sulfate (LDS) sample buffer and boiled for 2 min before being analyzed with SDS-PAGE and western blotting using anti-RhoA antibodies (Tchakal-Mesbahi et al., 2024). Quantification was performed using ImageJ software reflecting the relative amounts as a ratio of each protein band relative to the lane’s loading control. Ratio values represent three independent experiments.

### F-actin depolymerization assay

We plated LLC-AQP2 cells on 24 well plates and incubated them in normal DMEM for 2 days. Then, DMEM was replaced with HBSS and cells were incubated for 2 hours. After treatment with LVP/FK (10 nM/1 µM), DAPT (35 µM, 30 min), or the actin-depolymerizing drug latrunculin A (0.1 µM, 30 min), HBSS buffer was removed and cells were incubated in binding buffer containing phalloidin (20 mM KH_2_PO_4_, 10 mM PIPES, 5 mM EGTA, 2 mM MgCl_2,_ 4% PFA, 0.1% TritonX-100 and 250 nM rhodamine phalloidin (Sigma, St. Louis, MO) for 15 min at room temperature as previously described (Yui et al., 2012). Negative controls to estimate background autofluorescence were prepared using binding buffer lacking rhodamine-phalloidin. Cells treated with binding buffer were washed 4 times in PBS, incubated in 300 µl (in each well) methanol overnight at −20°C to extract bound rhodamine-phalloidin. The extracted rhodamine-phalloidin fluorescence was read using a Beckman DTX-880 multi-plate reader (excitation 535 nm, emission 595 nm). F-actin content values were expressed as relative fluorescence after subtraction of the negative-control values. Each fluorescence value was expressed as relative fluorescence unit (RFU)/well and data were analyzed using a two-tailed Student’s *t*-test (Yui et al., 2012).

### Kidney tissue slice preparation and immunostaining

To determine whether the effect of DAPT occurs in collecting duct PC *in situ*, AQP2 localization was studied in rat kidney tissue slices incubated *in vitro* as reported in our previous studies (Babicz et al., 2025; Cheung et al., 2019; Cheung et al., 2017; Li et al., 2011). All procedures described were reviewed and approved by the Massachusetts General Hospital (MGH) Subcommittee on Research Animal Care and were performed in accordance with the National Institutes of Health Guide for the Care and Use of Laboratory Animals. Briefly, Sprague-Dawley rats (250 - 300 g) were anesthetized using isoflurane inhalation. Kidneys were perfused with HBSS (110 mM NaCl, 5 mM KCl, 1.2 mM MgSO_4_, 1.8 mM CaCl_2_, 4 mM sodium acetate, 1 mM Na_3_ citrate, 6 mM D-glucose, 6 mM L-alanine, 1 mM NaH_2_PO_4_, 3 mM Na_2_HPO_4_, 25 mM NaHCO_3_, pH 7.4) equilibrated with 5%CO_2_, 95%O_2_ at 37°C. A razor blade was used to cut the kidneys into approximately 2-3 mm-thick slices, which were further cut into 0.5 mm slices with a Stadie-Riggs microtome (Thomas Scientific, Swedesboro, NJ). Slices were incubated at 37°C for 30 min in HBSS before treatment, followed by 15 min with AVP (20 nM) as a positive control, for 30 min with DAPT (35 µM) or DMSO (10 ul - a negative control). Slices were then fixed by immersion in 4% paraformaldehyde fixative at room temperature for 30 min and vials were stored in fixative overnight at 4°C. After rinsing 3 x10 min in 10 mM sodium phosphate buffer containing 0.9% NaCl, pH 7.4 (PBS), slices were stored in the same buffer plus 0.02% NaN_3_ at 4°C until sectioning. For immunostaining, the kidney slices were processed following our well-established procedures (Eaton et al., 2024). Tissues were incubated in PBS containing 30% sucrose overnight and these cryoprotected slices were mounted on a cutting block in OCT compound 4583 (Tissue-Tek; Miles Inc., Elkhart, IN). Tissues were cut into 5 μm sections using a Leica CM3050S cryostat (Leica, Buffalo Grove, IL), and attached to Superfrost Plus glass slides (Fisher Scientific, Pittsburgh, PA), rehydrated in PBS for 20 min, treated with 1% SDS in PBS for 5 min as an antigen retrieval step (Brown et al., 1996), and washed three times with PBS. Sections were then incubated 10 min in PBS containing 1% bovine serum albumin as a blocking step and incubated overnight at 4°C with a goat anti-AQP2 antibody (Santa Cruz) diluted in PBS (0.4 μg/ml). After incubation, sections were washed 3 x 5 min in PBS and secondary antibodies were applied: donkey anti-goat IgG conjugated to Alexa 488, and for double incubations, donkey anti-rabbit IgG conjugated to Cy3 or donkey anti-mouse IgG conjugated to Cy3 (all secondary antibodies were used at 7.5 μg/ml) respectively for 1 h at room temperature. After 3 washes with PBS, nuclear DNA was visualized by staining with DAPI, 20 µg/ml for 10min. The stained sections were washed 3 times with PBS then mounted on glass slides with Vectashield mounting medium. The mounted slides were examined using a Nikon 80i widefield microscope and a 40x plan-apochromatic lens. Images were captured using Nikon Elements software. The experiment was repeated 4 times using different animals each time.

### Quantification and analysis procedures

Western blot quantification was performed using ImageJ software reflecting the relative amounts as a ratio of each protein band relative to the lane’s loading control. For each protein (row), the same-sized frame was superimposed as a region of interest (ROI) on all bands across the lanes, analyzing one sample at a time. Each band was quantified, and a value was recorded. Using the same frame size, a background measurement was taken on a non-stained region either above or below each band within the same row. The ROIs for other protein rows, including loading controls, were measured in the same way. After all data for the bands and their corresponding backgrounds, along with those for the loading controls, were exported to a spreadsheet, calculations were performed. The inverted pixel density for each band and background was calculated using the formula 255 – X, where X is the value recorded by ImageJ. For each protein band and loading control, the net intensity was obtained by subtracting the inverted background value from the inverted band value. The final relative quantification was expressed as the ratio of the net intensity of the protein band to the net intensity of the corresponding loading control.

### Statistics

Statistics were performed with Prism software (GraphPad, La Jolla CA). Differences in means were then compared between control and treatment at each time point using the Student t-test (two tailed). Each experiment was repeated at least three times. Statistical significance was determined at a p value <0.05, but the actual p values are provided in the figures for each experimental condition.

## Results

### Notch inhibition stimulates AQP2 plasma membrane accumulation in cultured cells

LLCPK1 cells stably expressing AQP2 were treated with VP (10 nM, 15 min) and the Notch inhibitor DAPT (35 μM, 30 min). VP induced AQP2 plasma membrane accumulation in LLCPK1-AQP2 cells, consistent with our previous results (Babicz et al., 2025; Cheung et al., 2019; Tchakal-Mesbahi et al., 2024). We also found that DAPT treatment resulted in significant AQP2 membrane accumulation in renal epithelial cells, as shown by immunofluorescence staining (Fig. 1A and 1B; mean ± SEM, n=3). This concentration was chosen after examining LLCPK1-AQP2 cells treated with different concentrations of DAPT (5, 10, 25, 35, and 70 μM) for 30 min. Membrane accumulation of AQP2 was already maximal in cells treated with 35 μM DAPT, a dose at which no toxicity was detectable (Supplemental Fig.1) (Xu et al., 2021).

**Fig. 1.**
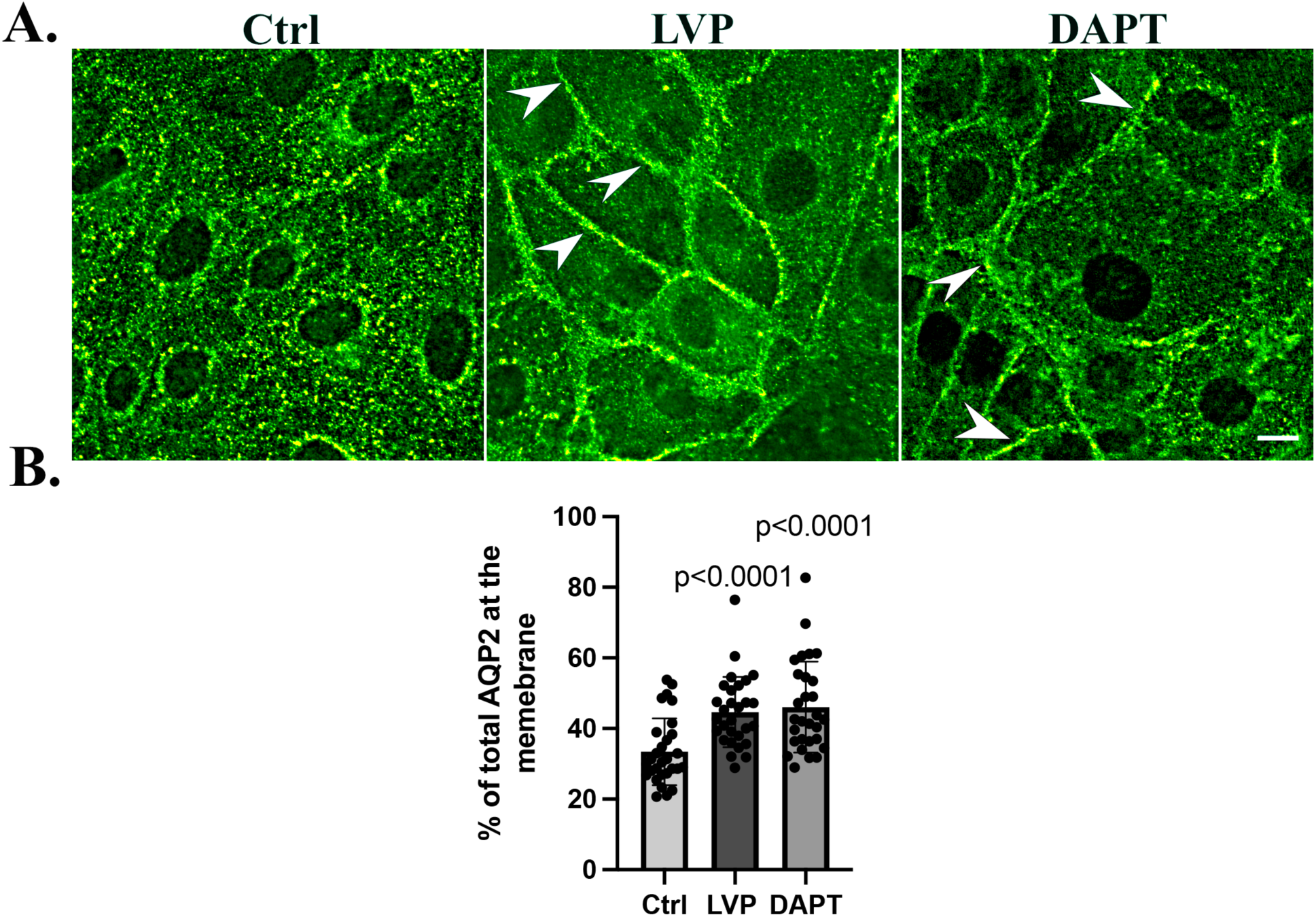
Notch inhibition induces membrane accumulation of AQP2 in LLCPK1-AQP2 cells. A. Immunofluorescence staining of AQP2 with anti-c-myc antibody in LLC-AQP2 cells treated with 35 μM DAPT for 30 min indicates a strong membrane accumulation of AQP2 (arrowheads, right panel) as also induced by LVP treatment (10 nM, 15 min; arrowheads, middle panel). B. Quantification showed that treatment with DAPT is indeed able to induce an increase in AQP2 in the membrane. Cells expected showed increased membrane accumulation of AQP2 in response to VP, to the same extent to DAPT. These results are representative of three independent experiments (n=3). Scale bar, 10 μM.

### Notch inhibition by DAPT decreases cleaved Notch 1 in renal epithelial cells

To confirm Notch signaling inhibition by DAPT in LLC-PK1–AQP2 cells, we examined cleaved Notch1 protein expression following DAPT treatment (35 μM, 30 min). Successful inhibition was confirmed by a decrease in cleaved Notch1 signal, which is otherwise increased when Notch signaling is active. LVP treatment showed no significant change in cleaved Notch1 levels compared to control (ns), whereas DAPT treatment significantly reduced cleaved Notch1 levels, confirming effective inhibition of Notch signaling (Fig. 2A and 2B).

**Fig 2.**
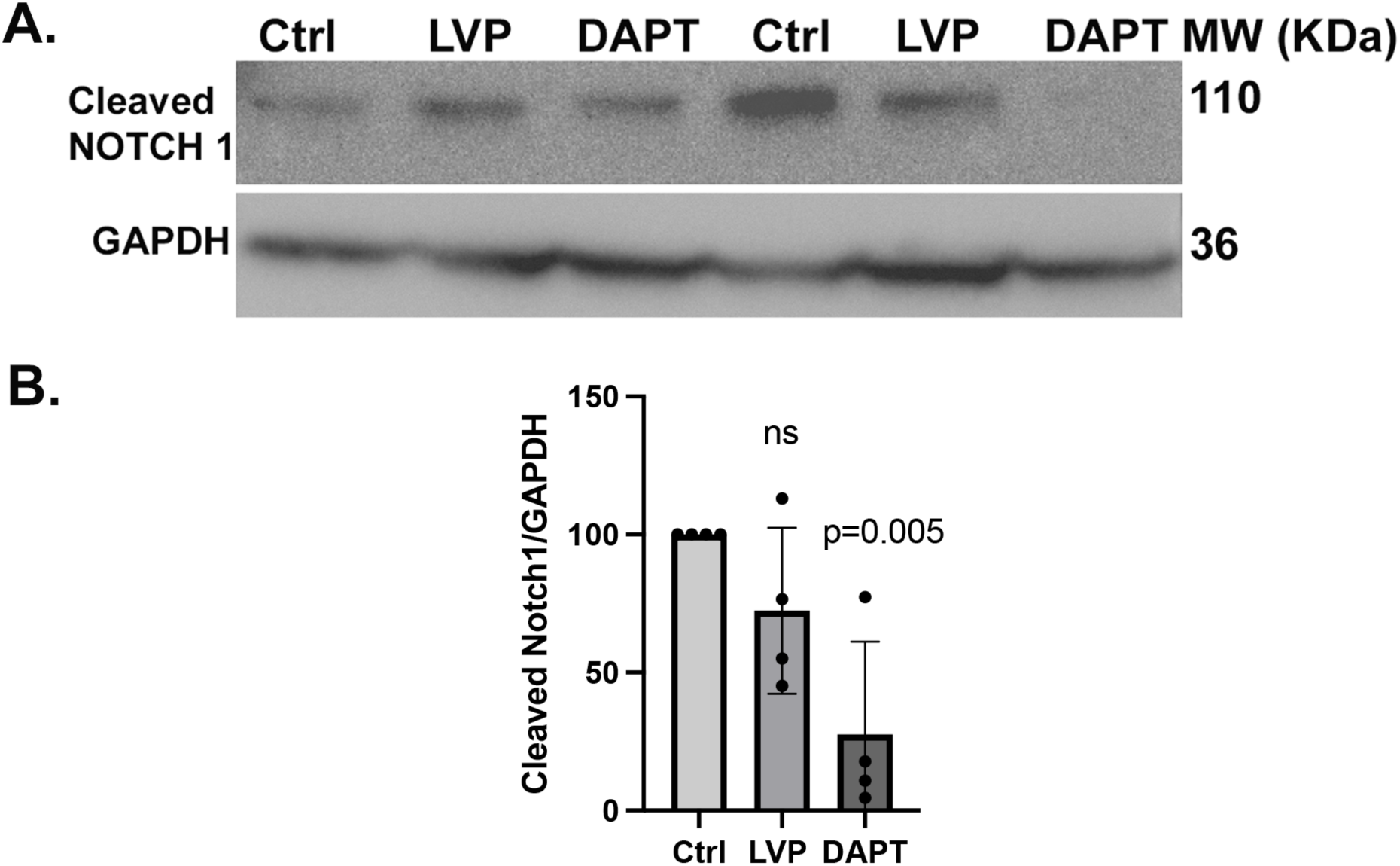
NOTCH inhibition in renal epithelial cells LLCPK1-AQP2 cells. **A.** Representative western blot showing cleaved Notch1 and GAPDH (loading control) protein levels in control (Ctrl), LVP-treated, and DAPT-treated cells. LVP treatment did not cause significant changes in cleaved Notch1 signal. Although some variability in cleaved Notch1 levels was observed between individual experiments, DAPT consistently inhibited Notch1 cleavage across 4 independent experiments. **B.** Bar graph of cleaved Notch1 normalized to GAPDH, expressed relative to control. LVP treatment showed no significant change compared to control (ns), whereas DAPT treatment significantly reduced cleaved Notch1 levels. Data are presented as mean ± SEM (n = 4 independent experiments).

### Notch signaling inhibition significantly reduces AQP2 endocytosis without significant changes to exocytosis in vitro

To understand how AQP2 accumulates at the plasma membrane, we studied both endocytosis and exocytosis pathways of AQP2 after Notch signaling inhibition. Using tetramethylrhodamine-conjugated transferrin as an endocytosis assay in LLCPK-AQP2 cells, we found that the cholesterol-depleting agent, MβCD efficiently blocked the endocytosis pathway as expected for this positive control. Under baseline conditions (control cells) tetramethylrhodamine (TMRh)-transferrin accumulated in the perinuclear region, as seen by immunofluorescence imaging. In contrast, MβCD and DAPT treatment resulted in significant transferrin accumulation on the cell membrane, i. e., it was not internalized by endocytosis. Quantification of the endocytosed TMRh-transferrin (Fig. 3A) confirmed that both MβCD and the Notch inhibitor, DAPT significantly decreased overall endocytosed TMRh-transferrin. The effect of MβCD was greater than that of DAPT, as expected for a drug that almost completely blocks all endocytotic pathways in cells. We also examined the effect of DAPT on endocytosis using the fluid phase marker Texas red dextran and found a significant 50% reduction of endocytosed dextran in LLCPK1-AQP2 cells, supporting the results using TMRh-transferrin, which is internalized by receptor mediated endocytosis (Fig. 3B).

**Fig 3.**
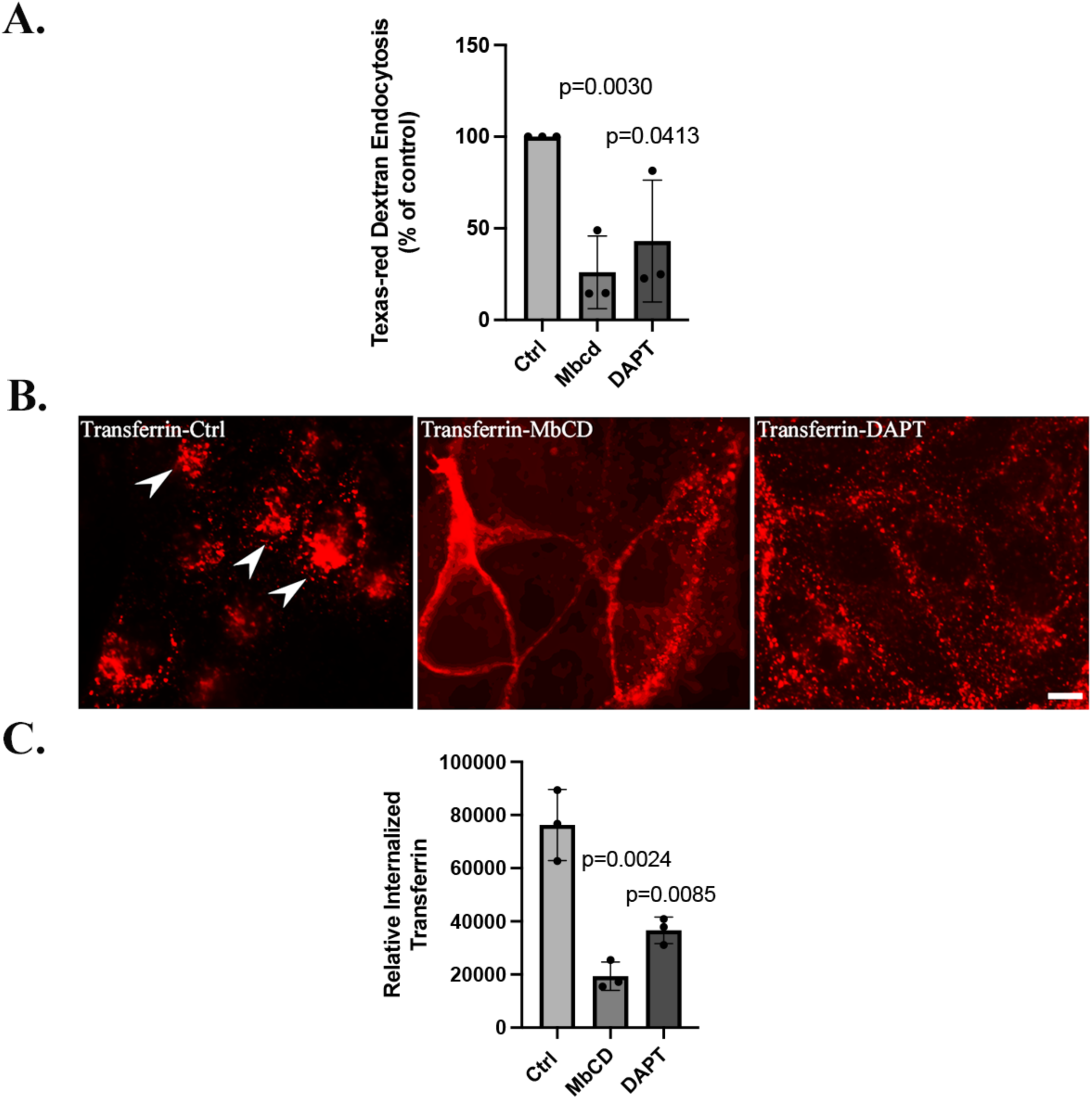
Notch inhibition decreases endocytosis in LLC-AQP2 cells. **A**. DAPT and MbCD (used as a positive control) reduced the endocytosis of Texas Red-tagged dextran in LLC-AQP2 cells. This result is an average of 3 independent experiments (n = 3). **B**. Rhodamine-tagged transferrin accumulates at the plasma membrane of LLC-AQP2 cells treated for 30 and 10 min with DAPT and MbCD respectively, in contrast to control cells (Ctrl), where transferrin is diffusely localized in the cytoplasm after endocytosis (mean ± SEM, n = 3. Bar is 10 μm. **C.** The quantification is the average of 3 independent experiments (mean ± SEM, n = 3). Bar is 10 μm.

**Fig. 4.**
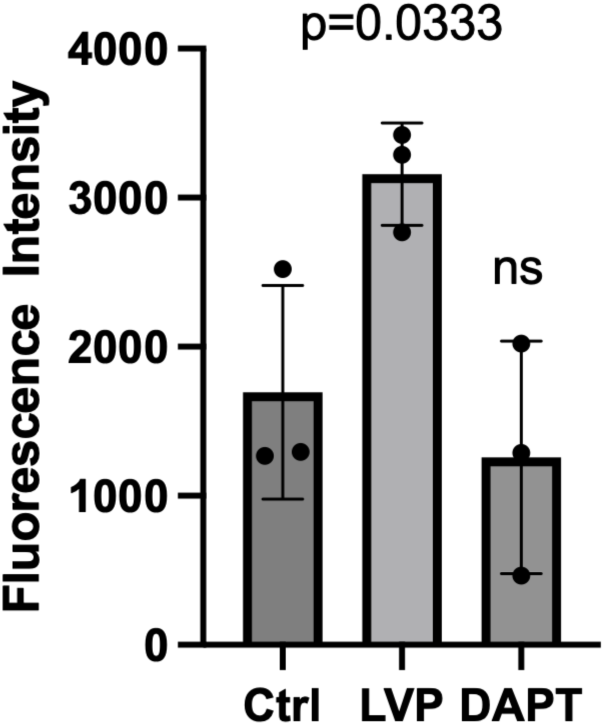
DAPT does not increase AQP2 exocytosis. Relative fluorescence of ssYFP secreted from LLC-AQP2-ssYFP cells into the culture medium was quantified under different conditions as a surrogate for AQP2 vesicle exocytosis. All experiments were repeated at least three times. Ctrl, control. LVP induce a significant increase in exocytosis as previously reported (Nunes et al., 2008), and DAPT treatment did not induce a significant increase in ssYFP intensity. Bar values represent means + SEM. Student’s t-test was performed to examine significance of treated groups vs. control (n = 3).

We then performed an exocytosis assay using LLCPK1-AQP2-ssYFP cells by measuring the surrogate marker of AQP2 exocytosis, soluble secreted YFP in the cell culture medium. A significant increase in the ssYFP exocytosis rate was found after LVP treatment as we previously described (Cheung et al., 2023; Lu et al., 2004). However, DAPT treatment did not induce any significant change in exocytosis as measured by ssYFP secretion from the cells (Fig. 34). We conclude that that Notch inhibition significantly reduces AQP2 endocytosis but without any significant effect on exocytosis in our cell culture systems. This endocytosis blockade would result in accumulation of constitutively recycling AQP2 on the plasma membrane, as we have reported for other drugs that block the endocytotic pathway (Cheung et al., 2019; Lu et al., 2004; Tchakal-Mesbahi et al., 2024).

### Notch inhibition causes AQP2 membrane accumulation independently of AQP2 phosphorylation at serine 256

AQP2 phosphorylation at serine 256 (S256) plays a critical role in regulating AQP2 trafficking. S256 phosphorylation alone is necessary and sufficient to cause AQP2 membrane accumulation upon VP signaling (Cheung et al., 2023). We used an AQP2 p-256 specific antibody to follow AQP2 phosphorylation after treatment with the Notch inhibitor DAPT. Western blots showed that LVP caused a significant increase in phosphorylation at S256, consistent with previous results (Katsura et al., 1997; Li et al., 2011; Moeller et al., 2010). In contrast, no significant difference in S256-AQP2 phosphorylation intensity was detectable after DAPT treatment (35 µM, 30 min) (Fig. 5). Additionally, we found that DAPT exposure also resulted in AQP2 membrane accumulation in our mutant cell line stably expressing an S256A mutation that prevents phosphorylation at this site. Consistent with prior data, LVP failed to induce membrane accumulation in these cells. Furthermore, treatment with the endocytosis blocker MβCD (Russo et al., 2006) also resulted in AQP2 accumulation on the plasma membrane of AQP2 S256A expressing cells (Fig. 5B). These findings show that Notch inhibition promotes AQP2 membrane accumulation independently of the S256 AQP2 phosphorylation pathway.

**Fig 5.**
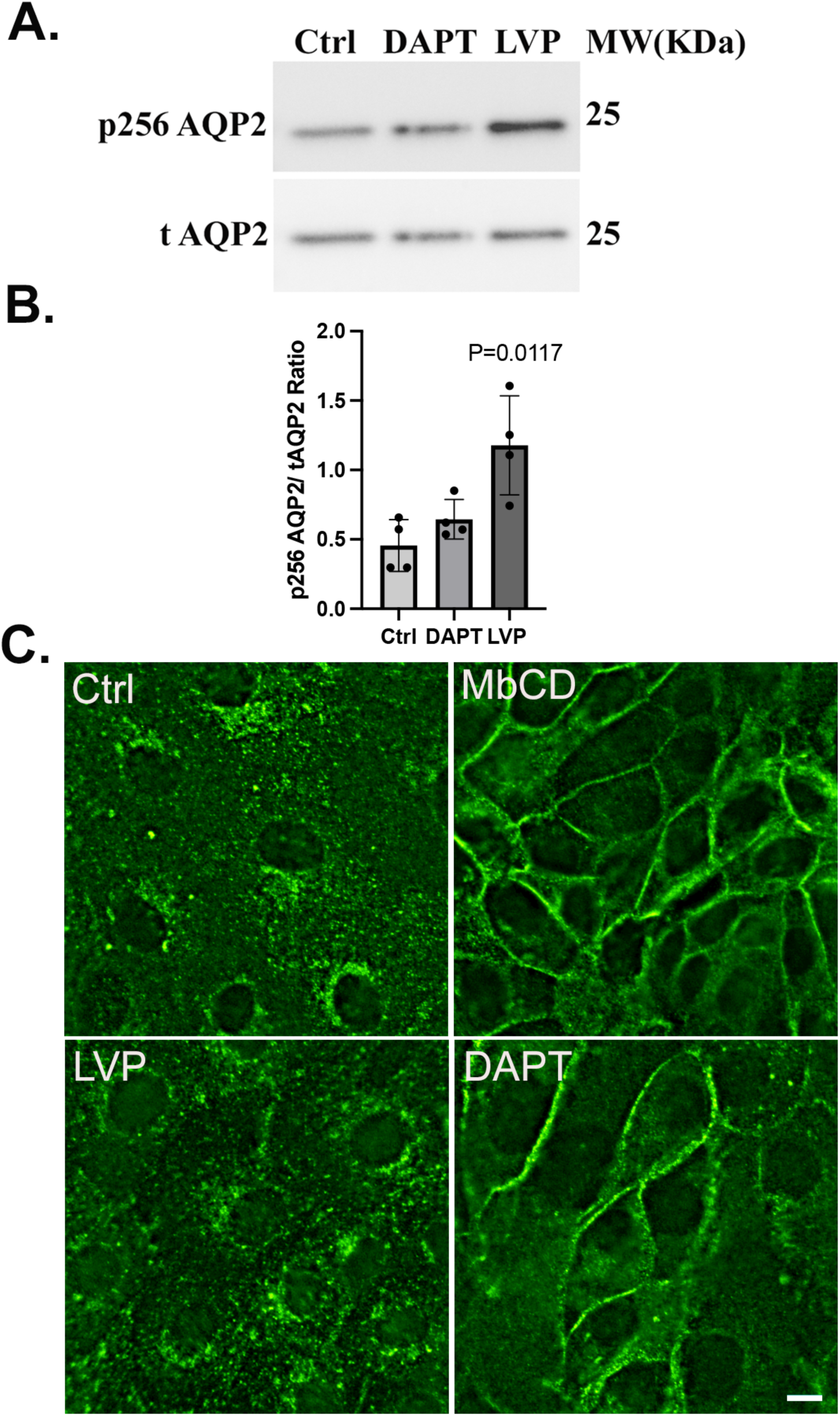
DAPT induces AQP2 membrane accumulation without phosphorylation at serine 256. **A**. Western Blot of total (t)AQP2 and pS256 indicates that DAPT does not induce AQP2 trafficking through the canonical, VP-dependent pathway because no significant increase in the phospho-S256 signal was detectable, in contrast to LVP treatment in 4 independent experiments. After detecting p256AQP2 with specific anti-p256AQP2 antibody, membranes were stripped and reblotted with tAQP2 antibody. **B**. Bar graph represents ratio of p256 AQP2 and tAQP2 band intensities of 4 independent experiments (mean ± SEM, n=4). Ratio analysis showed a consistent effect of LVP, whereas DAPT treatment showed a slightly higher ratio compared to control (Ctrl) cells with no significant effect on AQP2 phosphorylation. **C**. Membrane accumulation of AQP2 is induced by DAPT in cells expressing the S256A phosphorylation mutant. Methyl-b-cyclodextrin (MbCD) is a positive control to block endocytosis, resulting in AQP2 membrane accumulation in S256A-expressing cells; scale bar=10 μm.

### Notch inhibition does not induce changes in phosphorylation states of AQP2 at S269 and S261

To determine whether Notch inhibition affects AQP2 phosphorylation at additional regulatory sites, we examined the phosphorylation status of AQP2 at Ser269 (p269) and Ser261 (p261) following DAPT treatment. Western blot analysis of total AQP2 (tAQP2) and p269 AQP2 showed no significant change in the p269/tAQP2 ratio between control and DAPT-treated cells (Fig. 6A and 6B; mean ± SEM, n=3), indicating that DAPT does not induce AQP2 phosphorylation at Ser269. Similarly, analysis of p261 AQP2 relative to tAQP2 revealed no significant difference following DAPT treatment (Fig. 6C and 6D; mean ± SEM, n=4), demonstrating that Notch inhibition does not induce dephosphorylation of AQP2 at Ser261. These results are in contrast to the effect of VP on these two residues (Hoffert et al., 2008; Hoffert et al., 2006; Moeller et al., 2009; Rice et al., 2012).

**Fig. 6.**
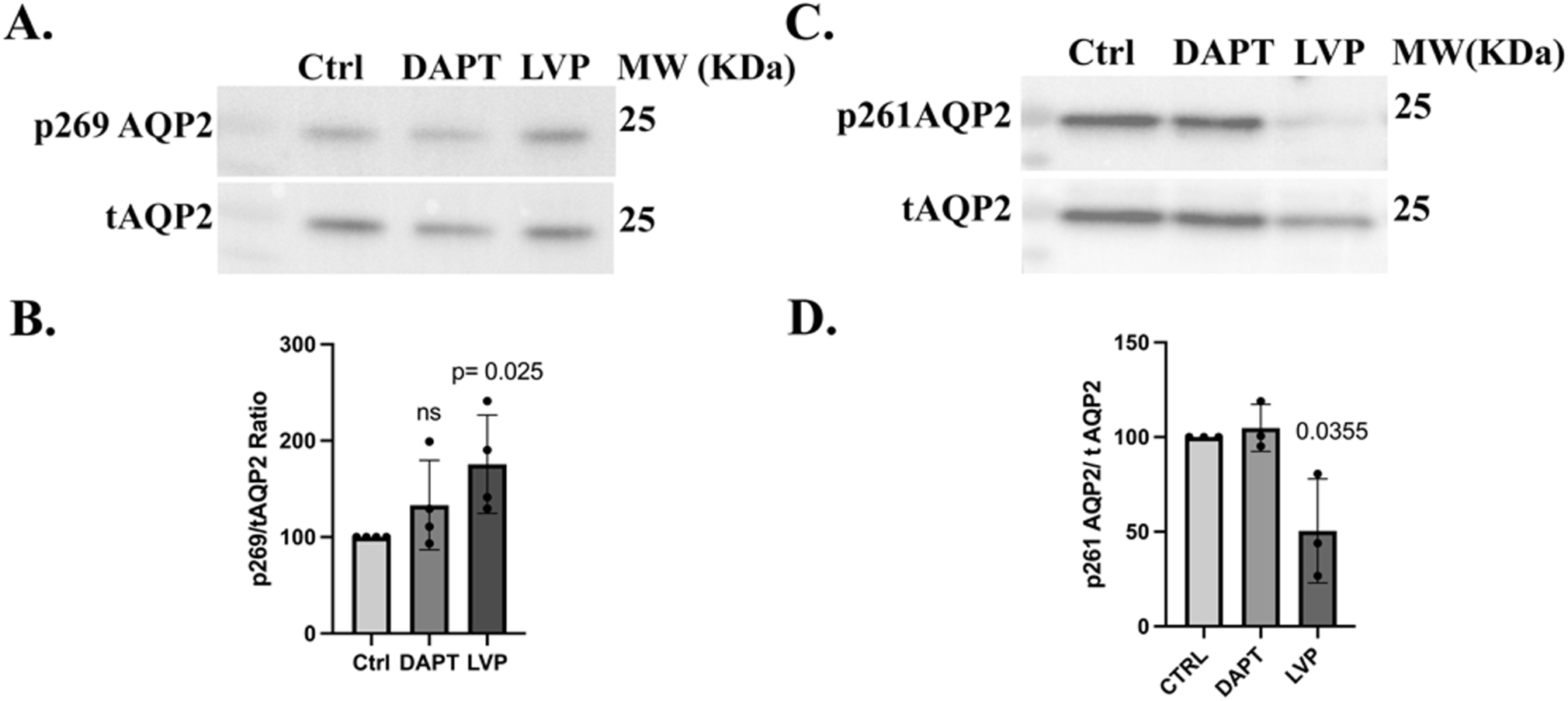
Notch inhibition does not affect the phosphorylation state of AQP2 at S269 and at S261. **A.** Western Blot of tAQP2 and p269 indicates that DAPT does not induce AQP2 phosphorylation at this residue. After detecting p269AQP2 with specific anti-p269 AQP2 antibody, membranes were stripped and reblotted with tAQP2 antibody. **B**. Bar graph represents ratio of p269 AQP2 and total AQP2 band intensities of 3 independent experiments (mean ±SEM, n=3). **C**. Western Blot of tAQP2 and p261 indicates that DAPT does not induce AQP2 dephosphorylation at S261. After detecting p261AQP2 with specific anti-p261 AQP2 antibody, membranes were stripped and reblotted with tAQP2 antibody. **D**. Bar graph represents ratio of p261 AQP2 and total AQP2 band intensities of 4 independent experiments (mean ± SEM, n=4).

### NOTCH inhibition induces AQP2 membrane accumulation and decreases cellular F-actin content

The actin cytoskeleton plays a key role in defining AQP2 dynamics and regulating its trafficking through polymerization and depolymerization. We applied LVP/FK (LVP 20 nM, FK 10 μM) and 35 μM DAPT to LLC-AQP2 cells for the indicated times, then subjected them to F-actin quantification. As a positive control, we used a depolymerizing agent, latrunculin (0.1 μM, 30 min) which caused a sharp 50% decrease in F-actin content (Fig. 7A). After LVP/FK treatment, the intracellular F-actin amount decreased significantly (about 20%) in LLC-AQP2 cells compared to untreated cells, and DAPT treatment resulted in a similar reduction in cellular F-actin content. To examine this process further, we used an assay that reveals the activation state of the small GTPase Rho A, which regulates actin polymerization. Compared to untreated cells, the amount of GTP-RhoA was strongly reduced after Notch inhibition, consistent with an attenuation of Rho activity (Fig. 7B and 7C). As expected, a large reduction in GTP-bound cellular RhoA was also observed in LLC-AQP2 cells treated with VP, which is known to inactivate RhoA activity. A reduction in RhoA activity is expected to reduce actin polymerization and is consistent with a reduction in total F-actin content of the cells after DAPT and VP treatment.

**Fig. 7.**
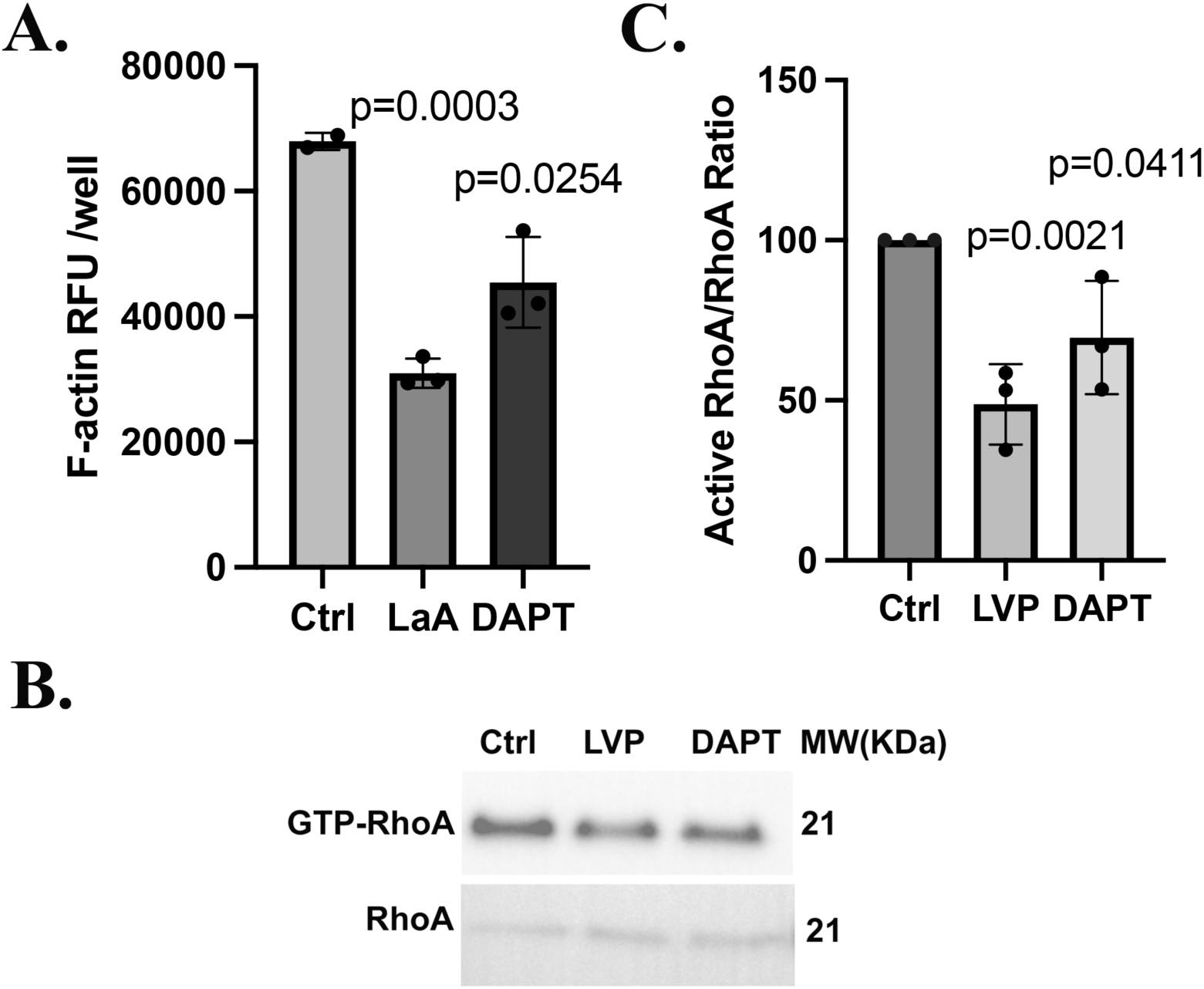
DAPT induces AQP2 membrane accumulation by inhibiting RhoA and depolymerizing F-actin. **A**. F-actin quantification assay using rhodamine-phalloidin binding shows that DAPT cause a significant 28% decrease in F-actin compared to controls. Latrunculin A, an established actin depolymerizing agent was used as a positive control and it induced a 50% decrease of cellular F-actin (mean ± SEM, n=3, p<0.01 and p<0.001). **B.** RhoA pulldown assay using GSTRBD beads that bind to active RhoA in LLC-AQP2 cells shows that DAPT reduces active RhoA in a similar way to vasopressin. D. Bar graph represents ratio of active RhoA and total RhoA band intensities of 3 independent experiments (mean ±SEM, n=3).

### Notch inhibition stimulates apical membrane accumulation of AQP2 in cultured rat kidney slices

To determine whether DAPT has a similar effect in kidney collecting duct principal cells in situ, we used rat kidney tissue slices as previously described in many of our previous studies (Babicz et al., 2025; Cheung et al., 2017; Li et al., 2011; Tchakal-Mesbahi et al., 2024). Both DAPT (35 µM for 30 mins) and VP (20 nM for 15 mins) resulted in a significant apical localization of AQP2 as detected by immunocytochemistry (Fig. 8). AQP2 had a diffuse intracellular localization in PCs from control, untreated tissues, whereas kidney slices incubated with DAPT for 30 min showed mostly a strong apical staining in collecting ducts from both the outer and inner medulla (Fig. 8). This pattern of staining was indistinguishable from that produced by 15 min of AVP treatment. Some basolateral staining was seen in PCs from the inner medulla, consistent with previous studies (Jeon et al., 2003; Nielsen et al., 1993; Tchakal-Mesbahi et al., 2024). These results show that the effect of the NOTCH inhibitor DAPT on AQP2 distribution is not restricted to our cell culture model but also occurs in collecting duct PC in situ.

**Fig. 8.**
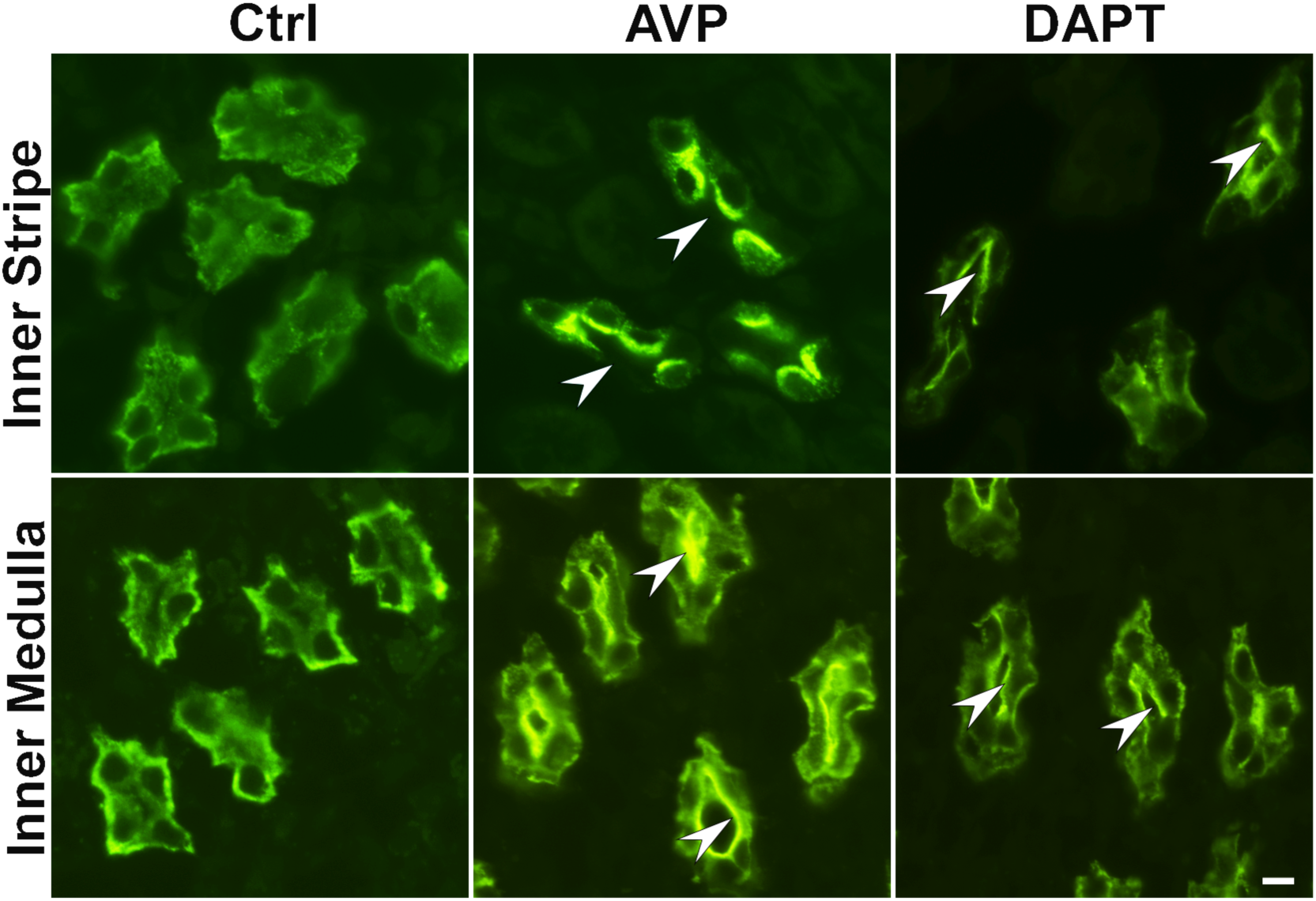
DAPT induces AQP2 membrane accumulation in kidney collecting duct principal cells. Using rat kidney slices in vitro as an experimental model, both AVP (20 nM; 15 min) and DAPT treatment (35 μM; 30 min) resulted in markedly increased apical staining in collecting duct PC. In the inner stripe of the outer medulla (upper panels), both treatments resulted in increased apical AQP2 accumulation compared to untreated tissues (Controls) in which diffuse cytoplasmic staining was predominant. Similarly, AVP stimulated apical AQP2 accumulation in the inner medulla, an effect that was reproduced by the Notch inhibitor DAPT (lower panels). Note that some basolateral membrane stainin is detectable in both control and stimulated conditions, as previously reported especially in the inner medulla (De Seigneux et al., 2007; Nielsen et al., 1993; Tchakal-Mesbahi et al., 2024; Yui et al., 2013). These images are representative of 3 independent experiments (n=3).

## Discussion

We show here that Notch signaling is involved in regulating AQP2 distribution and membrane accumulation both in cultured renal epithelial cells expressing exogenous AQP2, and in kidney principal cells in situ. This result contributes to our understanding of cellular signaling pathways that contribute to AQP2 recycling in the kidney, and that could ultimately suggest novel strategies to activate or inhibit AQP2 membrane accumulation in water balance disorders. Importantly, no cellular toxicity was observed with a similar concentration of DAPT that was reported in a previous study (40 μM) that resulted in the highest measurable inhibition of NOTCH signaling (Xu et al., 2021).

The small molecule (N-[(3,5-Difluorophenyl) acetyl]-L-alanyl-2-phenyl]glycine-1,1-dimethylethyl ester (DAPT) is a powerful tool to examine the Notch signaling pathway. It inhibits γ-secretase activity thereby preventing Notch receptor enzymatic hydrolysis, which blocks intracellular domain release and transcriptional activation in the Notch pathway (Hans et al., 2020; Yin et al., 2020). Importantly, DAPT is reported to have little or no effect on other cellular signaling pathways (Sun et al., 2019). Based on these considerations, we hypothesized that Notch signaling might regulate actin-dependent water channel trafficking in renal collecting duct cells. We propose that activation of the Notch pathway controls AQP2 membrane targeting by modulating RhoA-dependent cytoskeletal dynamics, thereby influencing water transport and renal concentrating ability.

Our findings show that inhibiting Notch with DAPT resulted in AQP2 accumulation at the membrane independently of its phosphorylation at S256. AQP2 traffics constitutively between cytosolic vesicles and the plasma membrane through continuous endocytosis and exocytosis in a very rapid and balanced way. Modification of this balanced recycling process by increasing or decreasing the relative rates of endocytosis or exocytosis results in greater of lesser accumulation of AQP2 at the cell surface (Brown, 2003; Lu et al., 2004; Sun et al., 2002). With this in mind, the focus of several studies on Notch and protein trafficking led us to investigate its role in AQP2 recycling. A systematic survey of endocytosis based on multiparametric image analysis has indicated that several pathways including Notch signaling regulate the endocytotic pathway (Collinet et al., 2010). In the kidney, Notch activation caused a loss of the slit diaphragm protein nephrin in podocytes by promoting dynamin-dependent endocytosis (Waters et al., 2012).

Interestingly, no change in AQP2 exocytosis was observed after NOTCH inhibition when we applied our ssYFP exocytosis assay. Previous studies have shown that AQP2 can accumulate at the membrane simply by inhibiting the endocytic pathway and that increased exocytosis is not required for AQP2 to accumulate in the plasma membrane (although increased exocytosis certainly occurs after VP stimulation as well as in other conditions). Using TMRh transferrin as an established assay to follow clathrin-mediated endocytosis, we showed that the effect of DAPT is a result of reduced endocytosis AQP2 without any detectable change in exocytosis (Yui et al., 2012).

As in several other instances in which a decrease in endocytosis is responsible for AQP2 membrane accumulation, phosphorylation of the S256 site is not required for AQP2 membrane localization after Notch inhibition. Indeed, we have previously reported similar findings after inhibition of the endocytic pathway or the depolymerization of actin (Liu et al., 2021). The actin cytoskeleton plays a key role in protein trafficking, and actin modulation through polymerization and depolymerization defines the cellular localization of many proteins in epithelial and non-epithelial cells (Goode et al., 2015; Ridley, 2006). Here, we found that inhibition of Notch signaling resulted in a significantly reduced amount of cytosolic F-actin in LLCPK1-AQP2 cells. The Notch inhibitor, DAPT significantly reduced the amount of active-GTP bound RhoA, which would explain the observed reduction in cellular F-actin. Actin dynamics play a key role in regulating AQP2 transport pathways including exocytosis, endocytosis and recycling (Mamuya et al., 2016; Tamma et al., 2003; Valenti et al., 2005). Indeed, it has been shown previously that actin depolymerization alone, without VP or any other stimulation, can shift AQP2 to the plasma membrane (Klussmann et al., 2001; Li et al., 2011; Lu et al., 2004).

Among the small GTPases, many studies have elucidated the major role of RhoA in reorganizing actin filaments by transitioning it between polymerized and depolymerized states to modulate protein trafficking and vesicle movements in the cell (Noda et al., 2004; Pochynyuk et al., 2007; Stirling et al., 2009). We and others have previously shown a direct effect of RhoA inactivation in inducing AQP2 membrane accumulation using statins in renal epithelial cells expressing AQP2 (Li et al., 2011; Procino et al., 2011). Our current findings align with these earlier data and now implicate Notch signaling in this process. There are many reasons to support Notch signaling in the regulation of actin polymerization (Belin de Chantemele et al., 2008; Tikka et al., 2012; Wang et al., 2011; Yui et al., 2012). Liu et al., showed that DAPT remodeled the F-actin cytoskeleton and inhibited the formation of lamellipodial cell migration, very likely by inhibiting non-canonical Notch signaling (Liu et al., 2019). Venkatesh et al demonstrated that Notch activates RhoA (Venkatesh et al., 2011) and the small GTPase RhoA can be activated by Notch signaling in certain cases (Belin de Chantemele et al., 2008). For instance, Notch3 functions as an upstream effector for RhoA/Rho kinase, and the lack of Notch 3 in tail arteries resulted in a significant decrease in active RhoA (Belin de Chantemele et al., 2008). Given these data, we propose that inhibition of Notch signaling inactivates RhoA to induce actin depolymerization, leading to low rate of endocytosis of AQP2, and subsequent plasma membrane accumulation.

Importantly, NOTCH inhibition mimicked the effect of VP on AQP2 localization in medullary collecting duct PC, stimulating both apical and/or basolateral plasma membrane accumulation in cells from different segments of the rat kidney collecting duct. This pattern of AQP2 polarity in response to VP has been described previously (De Seigneux et al., 2007; Nielsen et al., 1993) and it is not uncommon, possibly due to continuous AQP2 recycling through transcytosis (Yui et al., 2013), PC in the inner medulla in particular contains AQP2 on both basolateral and apical membrane, but the functional consequence of this bipolar distribution on collecting duct water permeability remains unclear.

Multiple studies have shown that although Notch is critical in individual development, the expression of Notch receptors and ligands is significantly decreased and kept at a lower expression and activation level in the mature organs compared with development (Surendran et al., 2010). In healthy adult glomeruli, Notch is almost absent, but Notch activation occurs in some kidney diseases including AKI (Kobayashi et al., 2008), and Notch inhibition rescues some of the symptoms (Kramer et al., 2016; Soni et al., 2019). Recent studies have also shown increased Notch receptor and ligand levels in various glomerular diseases (Murea et al., 2010; Niranjan et al., 2009; Walsh et al., 2008). It is unknown whether levels of Notch receptors and ligands are increased in NDI patients or mice, and this still requires further elucidation. Because of the low level of Notch in mature organs, Notch inhibition may actually be more beneficial in disease states such as NDI than potentially detrimental to normal structure and function. Given that up-regulation of the Notch signaling pathway is so common in multiple diseases, this pathway is also expected to be a potential target for combined therapy in various types of water balance disorders.

## Acknowledgements

This work was supported by the National Institutes of Health (NIH) DK096586-10A1 (D.B.), and a U2C/TL1 Harvard Kidney, Urology and Hematology Training Institute (HKUHTI) Network Grant (1TL1DK143273-01) (to A. M.) Additional support for the Program in Membrane Biology Microscopy Core comes from the Boston Area Diabetes and Endocrinology Research Center (P30DK135043) and the Massachusetts General Hospital (MGH) Center for the Study of Inflammatory Bowel Disease (P30DK43351). The Nikon AXR confocal microscope was purchased using an NIH S10 shared instrumentation grant 1S10D032211-01.

