## Supplemental data for "Inhibition of the Notch signaling pathway promotes AQP2 plasma membrane accumulation in renal epithelial cells by depolymerizing actin and reducing endocytosis"

**Supplemental table: List of Antibodies**

| Antigen | Manufacturer | Catalog | References | Dilution (WB) |
|---|---|---|---|---|
| RhoA | Cell Signaling | 2117T | PMID:37730597 | 1:1000 |
| GAPDH | Applied Biosystems | 4300 | PMID: 32434945 | 1:10000 |
| Cleaved Notch 1 | Cell Signaling | 4147 | PMID: 40190020 | 1:1000 |
| P256 AQP2 | Affinity | 3673 | PMID: 15509592 | 1:1000 |
| P269 AQP2 | Cell Signaling | 3674 | PMID: 15509592 | 1:1000 |
| P261 AQP2 | Rockland antibodies | 612-401 D08 | PMID: 7508187 | 1:1000 |
| AQP2 | Santa Cruz | 9882 | PMID: 22218592 | 1:1000 |

**Supplemental Figure 1:**

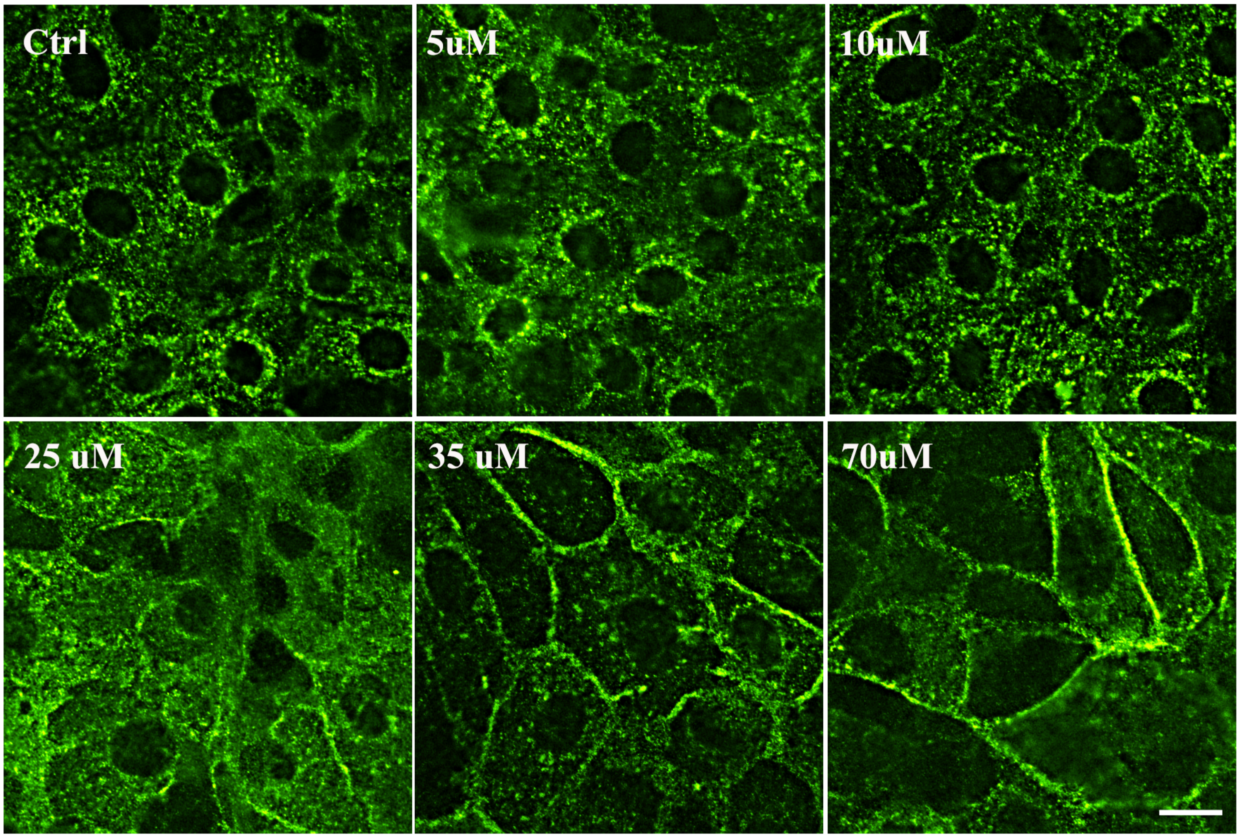


**Supplementary Figure 1. DAPT induces membrane accumulation of AQP2 in a dose-dependent manner in LLC-AQP2 cells**. Immunofluorescence staining of AQP2 with anti-c-myc antibody in LLCAQP2 cells treated with DAPT at various concentrations (5, 10, 25, 35 and 70) μM for 30 min. Membrane accumulation of AQP2 was detectable in cells treated with 25 μM DAPT and peaked at 35 μM DAPT. Scale bar = 10 μm.
